# Stem Cell Microarrays for Assessing Growth Factor Signaling in Engineered Glycan Microenvironments

**DOI:** 10.1101/2021.06.19.448747

**Authors:** Austen L. Michalak, Greg W. Trieger, Kelsey A. Trieger, Kamil Godula

## Abstract

Extracellular glycans, such as glycosaminoglycans (GAGs), provide an essential regulatory component during the development and maintenance of tissues. GAGs, which harbor binding sites for a range of growth factors and other morphogens, help establish gradients of these molecules in the extracellular matrix (ECM) and promote the formation of active signaling complexes when presented at the cell surface. As such, GAGs have been pursued as biologically active components for the development of biomaterials for cell-based regenerative therapies. However, their structural complexity and compositional heterogeneity make establishing structure-function relationships for this class of glycans difficult. Here, we describe a stem cell array platform, in which GAG polysaccharides are conjugated to adhesion proteins and introduced into a polyacrylamide hydrogel network to directly measure their contributions to the activation of growth factor signaling pathways in cells. With the recent emergence of powerful synthetic and recombinant technologies to produce well-defined GAG structures, a platform for analyzing both growth factor binding and signaling in response to the presence of these biomolecules will provide a powerful tool for integrating glycans into biomaterials to advance their biological properties and applications.

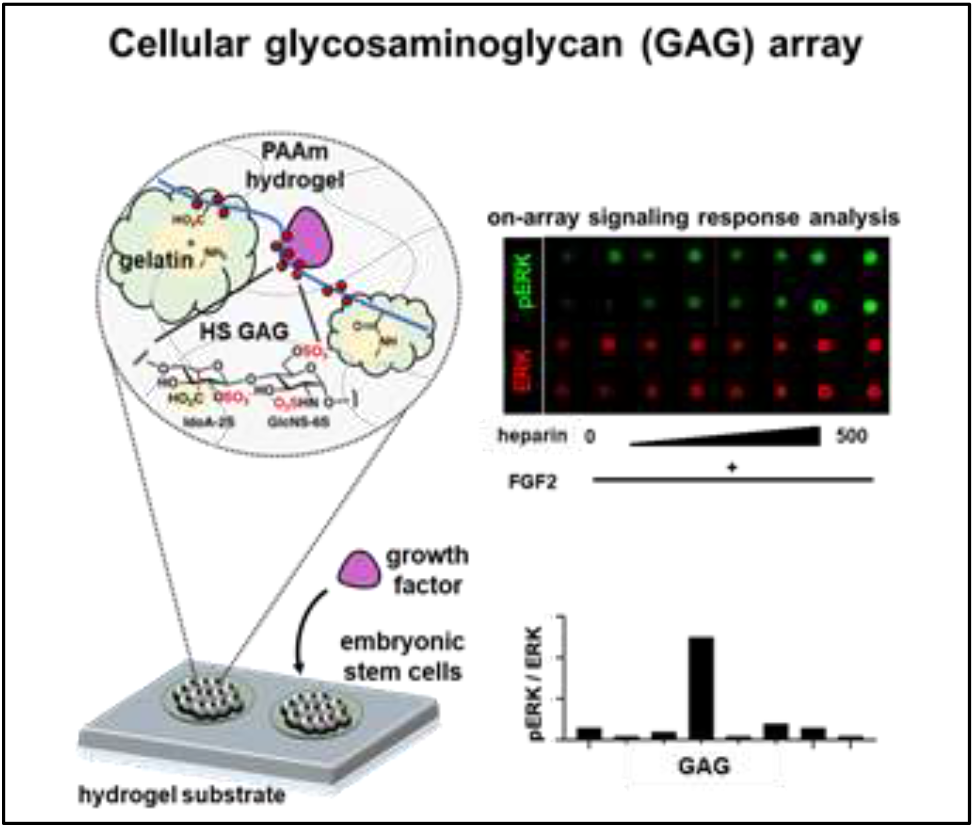

The present study describes the integration of glycosaminoglycan-protein conjugates into a hydrogel-supported stem cell microarray platform to analyze the activity of extracellular glycans in growth factor signaling. Such platforms can enable rapid development and optimization of functional glycomaterials for stem cell-based regenerative therapies.

## Introduction

The development of biologically active materials that support cell adhesion and proliferation, while also providing signaling cues to guide cellular differentiation, has enabled the translation of the regenerative capacity of stem cells into clinical applications.^[1,2]^ The integration of various components of the native extracellular matrix into hydrogels has emerged as a major strategy for generating responsive materials for organoid and tissue engineering.^[3,4]^ Comprised of hydrated synthetic or biological polymer networks, hydrogels are commonly decorated with peptides or proteins for cell adhesion and supplemented with signaling molecules, such as growth factors (GFs), to promote signaling and differentiation toward desirable cell types.^[5,6,7]^

Stem cell arrays, which allow for high-throughput analysis of cellular responses to their environment and culture conditions, have enabled the discovery and optimization of new biomaterials for cell-based applications.^[8]^ Such platforms have been particularly useful for examining the ability of various protein components of the ECM to enhance cell interactions and functions when introduced into hydrogels.^[9,10]^ Extracellular glycans, which also provide important biological functions in the ECM but are difficult to access in pure form synthetically or through isolation, have been comparatively less explored as components for biomaterials.^[11,12]^ For example, extracellular heparan sulfate (HS) polysaccharides, which belong to the family of glycosaminoglycans (**Figure 1**), are essential regulators of GF signaling and are being pursued as biologically active components of hydrogels for stem cell culture and tissue engineering.^[13]^ HS polysaccharides comprise chains of alternating *N*-acetylglucosamine and glucuronic acid residues, which undergo sequential enzymatic modifications to introduce *N*-sulfation and to partially epimerize GlcA into iduronic acid (IdoA).^[14]^ Additional *O*-sulfation is then introduced to produce sulfated domains harboring protein binding motifs. The compositional complexity of HS has made systematic structure-function analysis needed for their integration into biomaterials challenging. Recent advances in chemical^[15,16]^ and chemoenzymatic^[17,18]^ HS oligosaccharide synthesis as well as genetic engineering^[19]^ of HS biosynthetic pathways have produced increasingly large numbers of chemically well-defined HS structures available for examination in the context of biomaterial design.

**Figure 1.**
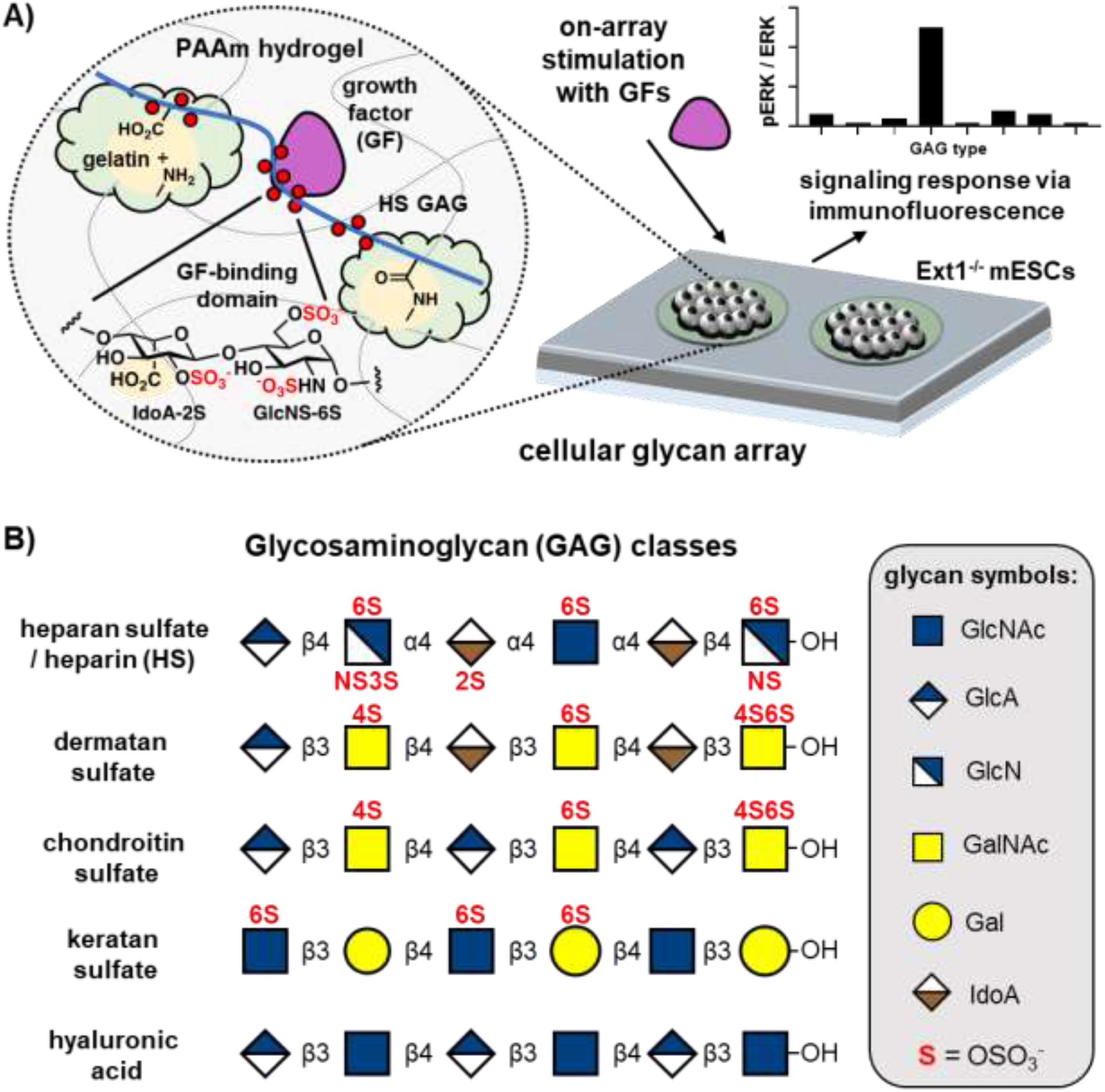
A cellular microarray approach to interrogate ECM-mediated cell signaling. A) Gelatin hydrogels on a non-biofouling acrylamide scaffold support ESC culture in the presence of highly sulfated crosslinked GAGs which direct cellular signaling. The array supports GF stimulation, which subsequently is assayed for cellular signaling using immunofluorescence. B) Structure and sulfation pattern of major GAG classes, with typical sulfation patterns denoted in red.

Arrays comprising isolated or synthetic HS structures immobilized on glass surfaces are routinely used to profile the specificity of HS-binding proteins.^20^ Developing of platforms to enable multiplexed, on-array analysis of HS-dependent cellular signaling could significantly streamline the discovery of biomaterials that capitalize on the regulatory functions of ECM glycans. An early example of arrays being used to evaluate the effects of HS structures on cellular responses came from Linhardt and co-workers, who studied proliferation of hydrogel encapsulated non-adherent Ba/F3 cells in the presence of chemically defined HS polysaccharides and fibroblast growth factors (FGFs).^[21]^ The cell-laden hydrogel droplets were printed on glass and exposed to combinations of HS and FGFs as soluble media supplements. Turnbull and his co-workers were able to directly observe activation of the mitogen activated protein kinase (MAPK) signaling pathway after FGF2 stimulation in Swiss 3T3 cells grown on arrays of oligosaccharides derived by partial heparin digestion. The cells were grown as a monolayer on HS oligosaccharides of increasing length (degree of polymerization, DP = 2-18) spotted and covalently immobilized on amine-functionalized glass via reductive amination. ^[22]^ MAPK activation was quantified by immunostaining for phosphorylation of Erk1/2 kinases and the magnitude of the observed signal scaled with oligosaccharide length.

To fully harness the multiplexing potential of these the array platform, strategies are needed to present HS structures to progenitor cells in a spatially isolated, yet addressable, format. Here, we present a method for the generation of hydrogel-based GAG microarrays for analysis of growth factor-mediated signaling in murine embryonic stem cells (ESCs). By arraying HS-protein conjugates on polyacrylamide hydrogels, we were able to generate stable ECM-mimetic microenvironments with the capacity to bind FGF2 and influence ESC signaling. The selectivity of FGF2 binding and activity in these cellular microenvironments was defined by the chemical composition of the HS polysaccharides.

## Results

To develop an array for assessing stem cell signaling responses to engineered glycan ECM environments, we sought to present the glycans together with cell adhesion factors in microscopic islands separated by a non-adhesive surface. This would enable multiplexed analysis of a range of glycan structures while minimizing cellular crosstalk. After screening several common surface passivation strategies used for array construction (**Figure S1**), we found that a thin poly(acrylamide) hydrogel deposited on glass according to a method by Brafman *et al.*^[23]^ and spotted with a solution of gelatin (500 μg/mL) in PBS best supported ESC growth in well-separated colonies over 6 days in culture. The gelatin, a commonly used substrate for murine ESC culture, was loaded into the hydrogel in its dehydrated form, which allowed the protein to enter and become entrapped within the crosslinked polymer network.

To test whether GAG polysaccharides may similarly be arrayed and retained within the hydrogel, the dry acrylamide substrates were spotted with gelatin solutions (500 μg/mL) in PBS buffer (10% glycerol, 0.003% triton X-100) supplemented with increasing concentrations (50-750 μg/mL) of heparin (12 kDa) as a model HS glycan (**Figure 2**). Anticipating that the polysaccharide may diffuse out of the hydrogel network under cell culture conditions, we also included conditions where the heparin was crosslinked via its carboxylic acid groups activated in the form of *N*-hydroxysuccinimide (NHS) esters to the solvent exposed lysine residues in gelatin (**Figure 1A**). The heparin was activated by treatment with NHS in HEPES buffer (100 mM, pH = 7.4) in the presence of the coupling reagent, 1-ethyl-3-(3-dimethylaminopropyl)carbodiimide (EDC), at 4°C for 18 hours and purified by size exclusion on a PD10 column to remove small molecule reagents and byproducts. We targeted low levels of crosslinking (∼ 15% carboxylic acid crosslinks per chain) to promote the retention of the GAG in the hydrogel network without compromising its GF-binding ability.

**Figure 2.**
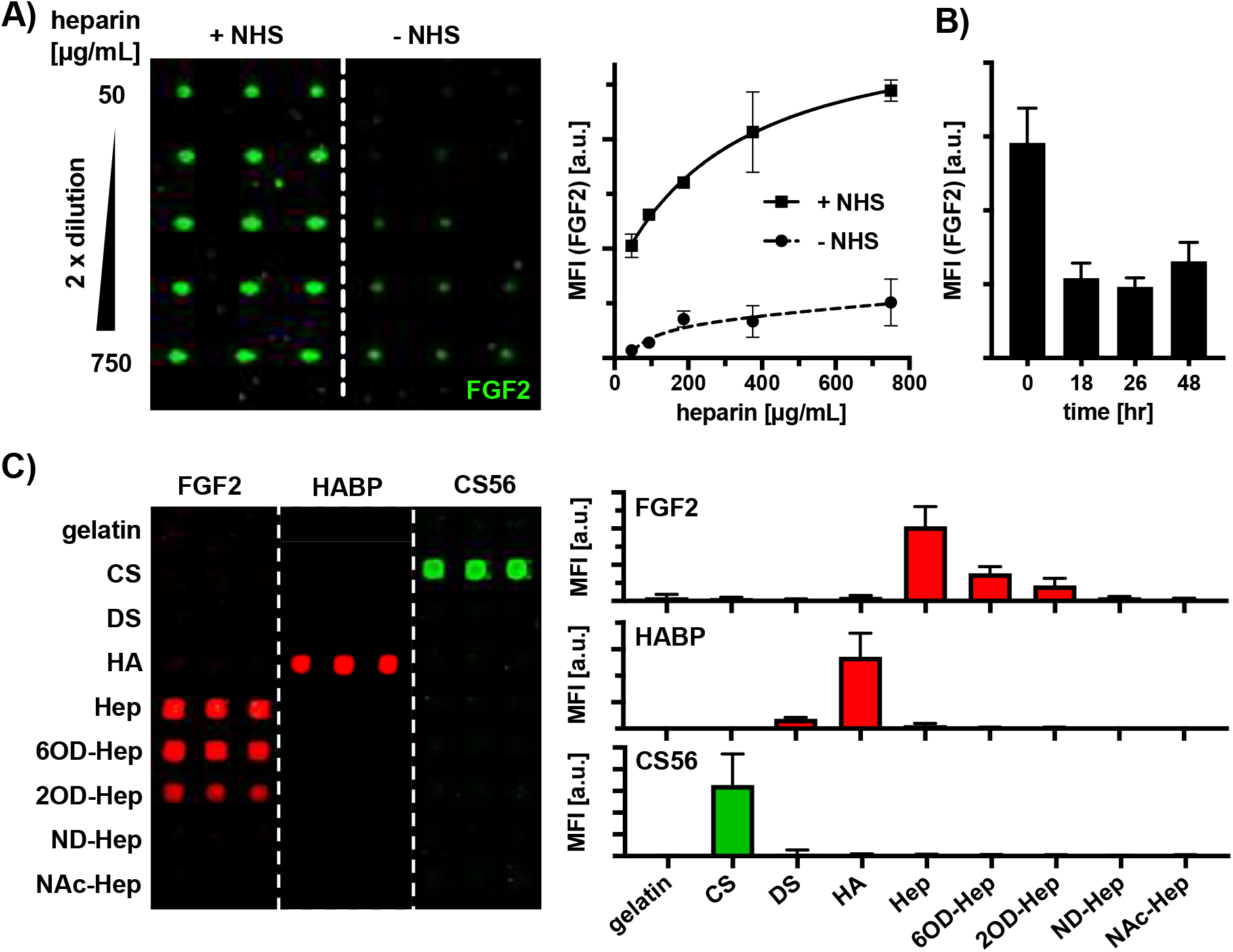
Generation and characterization of hydrogel glycosaminoglycan arrays for stem cell culture. A) Covalently linked heparin shows superior FGF2 binding capacity compared to non-covalently immobilized heparin on the array surface. Slides were washed for 3 hours prior to staining with fluorescently labeled FGF2 and imaging. B) FGF2 is retained on the surface of the array for 48 hours under cellular culture conditions at 37°C, 5% CO_2_. C) GAG molecular recognition and selectivity was assessed on the array using the FGF2, HABP, and CS56 to selectively bind heparin, hyaluronic acid, and CS, respectively. FGF2 binding to heparin showed the greatest preference for fully sulfated heparin, followed by 2O and 6O-sulfated sulfated heparin, respectively.

The resulting arrays were washed for 3 hours and probed with the heparin-binding FGF2 protein labeled with AlexaFluor 647 (FGF2-AF647) to detect the immobilized heparin and to assess the effect of crosslinking on its retention in the hydrogel (**Figure 2A**). Both immobilization strategies resulted in a dose-dependent FGF2 binding, with the NHS crosslinked heparin providing ∼ 5- to 10-fold higher signal. To further test the stability of the arrays, acrylamide substrates spotted with the NHS-activated heparin (500 μg/mL) in gelatin (500 μg/mL) were subjected to ESC culture conditions for 48 hours. Over this time, the arrays were probed with FGF2-AF647 to assess heparin retention (**Figure 2B**) After an initial decrease in binding activity during the first 18 hours, the heparin arrays remained stable with ∼ 40% of FGF2 binding activity being retained after 48 hours.

With a suitable method for heparin immobilization on the hydrogel substrates in hand, we aimed to test that the protein binding specificities of the crosslinked GAG structures within the hydrogel matrix are preserved (**Figure 2C**). Using the NHS-crosslinking strategy, we arrayed a panel of CS, DS, and HA GAGs as well as heparin polysaccharides chemically treated to selectively remove their 6-*O*-, 2-*O*-, and *N-*sulfates (6OD-Hep, 2OD-Hep, and ND-Hep, respectively). *N*-desulfated heparin, in which the exposed amino groups were capped as acetamides (NAc-Hep) to better represent native HS structures, was also included. The array was then probed with FGF2, a CS-specific antibody (CS56), and the hyaluronic acid binding protein (HABP). As shown in **Figure 2C**, FGF2 bound most strongly to the fully sulfated heparin. Removal of 6-*O*-, 2-*O*- and *N*-sulfates resulted in progressive loss of activity, which is in agreement with the known requirements of 2-*O*- and *N*-sulfation for FGF2 binding to HS.^[24]^ Likewise, CS56 and HABP proteins exhibited high specificity for CS and HA, respectively.

Having confirmed that NHS-crosslinking to the gelatin matrix enhances to stability of the GAG displays without altering the protein binding specificity of the polysaccharides, we set to evaluate the ability of these arrays to support ESC culture. For our cell model, we chose murine ESCs lacking the expression of *Exostosin* 1 (*Ext1*), which is a glycosyl transferase responsible for the assembly of HS chains.^[25]^ In the absence of this enzyme, the *Ext1^−/−^* ESCs lack cell surface HS structures and are unable to engage a range of HS-dependent GFs, ^[26]^ including FGF2. As such, these mutant ECS are ideally suited to isolate the effects of the arrayed GAGs on FGF2 signaling from those of endogenous HS structures.

The envisioned on-array GF signaling assay would require that the cells formed near-confluent monolayer colonies on the printed heparin-gelatin spots after at least 2 days in culture. In order to suppress endogenous GF production and establish signaling activity baseline, the last 24 hours should be carried out under serum-free conditions. To optimize cell density and colony growth on the array, we seeded increasing number of *Ext1^−/−^* ESCs on the substrates and grew them for 24 hours in embryonic culture media supplemented with leukemia inhibitor factor (LIF) and fetal bovine serum (FBS). The arrays were then washed, and the remaining bound cells were cultured for additional 24 hours in the absence of serum (**Figure 3A**). While seeding the stem cells too sparsely (20,000 cells/cm^2^) resulted in slow growth and irregular colony formation, too high seeding density (100,000 cells/cm^2^) led to rapid proliferation resulting in spot overgrowth and cell detachment. The intermediary seeding density (40,000 cells/cm^2^) produced consistent monolayers of *Ext1^−/−^* ESCs (**Figure 3A**), which retained high levels of expression of the embryonic marker, Oct4, and showed no obvious signs of differentiation (via the neural marker, Nestin**, Figure 3B**). The optimized seeding conditions were further tested in the presence of immobilized heparin printed at 500 μg/mL concentration and under serum-free starvation conditions to ensure no negative effects of these conditions on cell adhesion and growth (**Figure 3C**).

**Figure 3.**
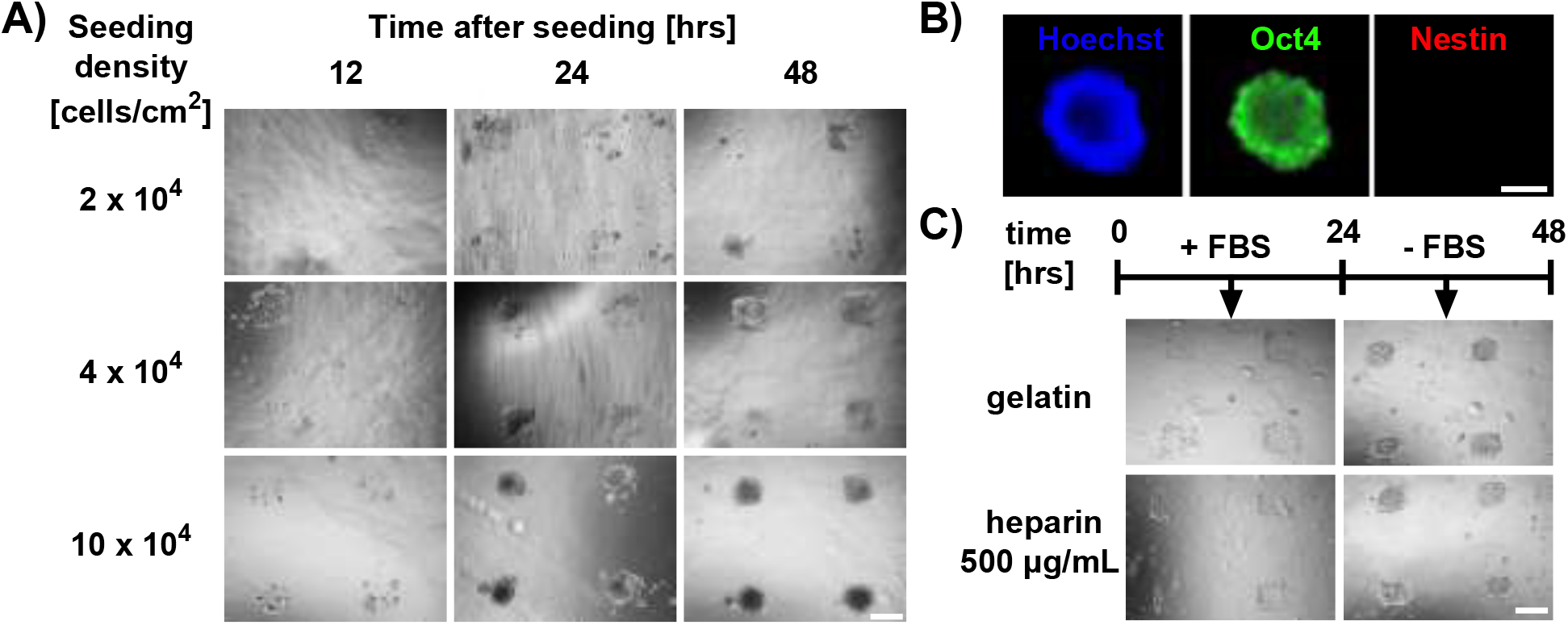
Establishment and optimization of microarray platform for stable cell culture of *Ext1^−/−^* ESCs. A) *Ext1^−/−^* ESCs seeded at densities of 2 ×10^4^, 4 ×10^4^, 10 ×10^4^ cells/cm^2^ on gelatin spots (0.5 mg/mL) arrayed on acrylamide substrates were assessed for growth and colony morphology over 48 hr period by optical microscopy (scale bar = 500 μm). B) After 48 hours on array culture in embryonic media containing LIF, the *Ext1^−/−^* ESCs retained high levels of pluripotency (Oct4) without significant spontaneous differentiation (Nestin). (scale bar = 200 μm. C) Immobilized heparin (500 μg/mL) does not significantly alter *Ext1^−/−^* ESCs adhesion, growth and colony formation on gelatin arrays. Cells were seeded at 4 ×10^4^ cells/cm^2^ and cultured for 48 hours, with the last 24 hours under serum-free conditions. (scale bar = 500 μm)

To establish whether the array format is suitable for directly assessing changes in stem cell signaling in the engineered glycan microenvironments, we chose to examine the activation of the MAPK pathway in response to stimulation with exogenous FGF2. The requirement for HS in the formation of a signaling complex between FGF2 and its receptor, FGFR, has been well established and the signaling response is accompanied by well-characterized changes in the phosphorylation status of downstream kinases (i.e., Extracellular regulated kinase 1 and 2, Erk1/2).^[27,28,29]^

For the on-array FGF2 signaling assay, *Ext1^−/−^* ESCs (40,000 cells/cm^2^) were seeded on spots printed with gelatin (500 μg/mL) with or without NHS-crosslinked heparin (500 μg/mL) and grown for 48 hours under the optimized embryonic culture and starvation conditions (**Figure 4A**). The cells were then placed in a fresh serum-free media containing FGF2 (0.5 ng/mL) and stimulated for 15 min at 37 °C. The cells were fixed, permeabilized, and immunoassayed for MAPK activity using antibodies against Erk1/2 proteins and their phosphorylated forms (pErk1/2). Fluorescence from the arrayed cells was detected using a microarrays scanner (**Figure 4B**) and validated via fluorescence microscopy (**Figure 4C**). The ratios of fluorescent signals corresponding to the phosphorylated-ERK1/2 (pERK, green) and total ERK1/2 (ERK, red) proteins was used to quantify the signaling response (**Figure 4B**). We used soluble heparin (s-Hep, 5 μg/mL), which is known to restore MAPK activity in *Ext1^−/−^* ESCs,^[19]^ as a positive control and a benchmark in our assay. While ERK1/2 protein levels were similar across all conditions, only *Ext1^−/−^* ESCs stimulated with FGF2 in the presence of immobilized or soluble heparin showed significant increase in Erk1/2 phosphorylation (**Figure 4B**). We performed fluorescent microscopy imaging **(Figure 4C)** and image J analysis (**Figure S2**) to confirm the co-localization of the pERK and ERK signals and to validate our quantification scheme based on signal detection via microarray scanner. We observed somewhat lower levels of MAPK activity on the arrayed heparin compared to its soluble form (**Figure 4B**). This may be due to the a more limited accessibility of the immobilized heparin to only a subset of FGFRs localized to the point of cell contact with the array. FGF2 stimulation of *Ext1^−/−^* ESCs grown on gelatin spots containing increasing amounts of immobilized heparin (i-Hep, 0-500 μg/mL) showed a heparin dose-responsive ERK1/2 phosphorylation (**Figure 4D**).

**Figure 4.**
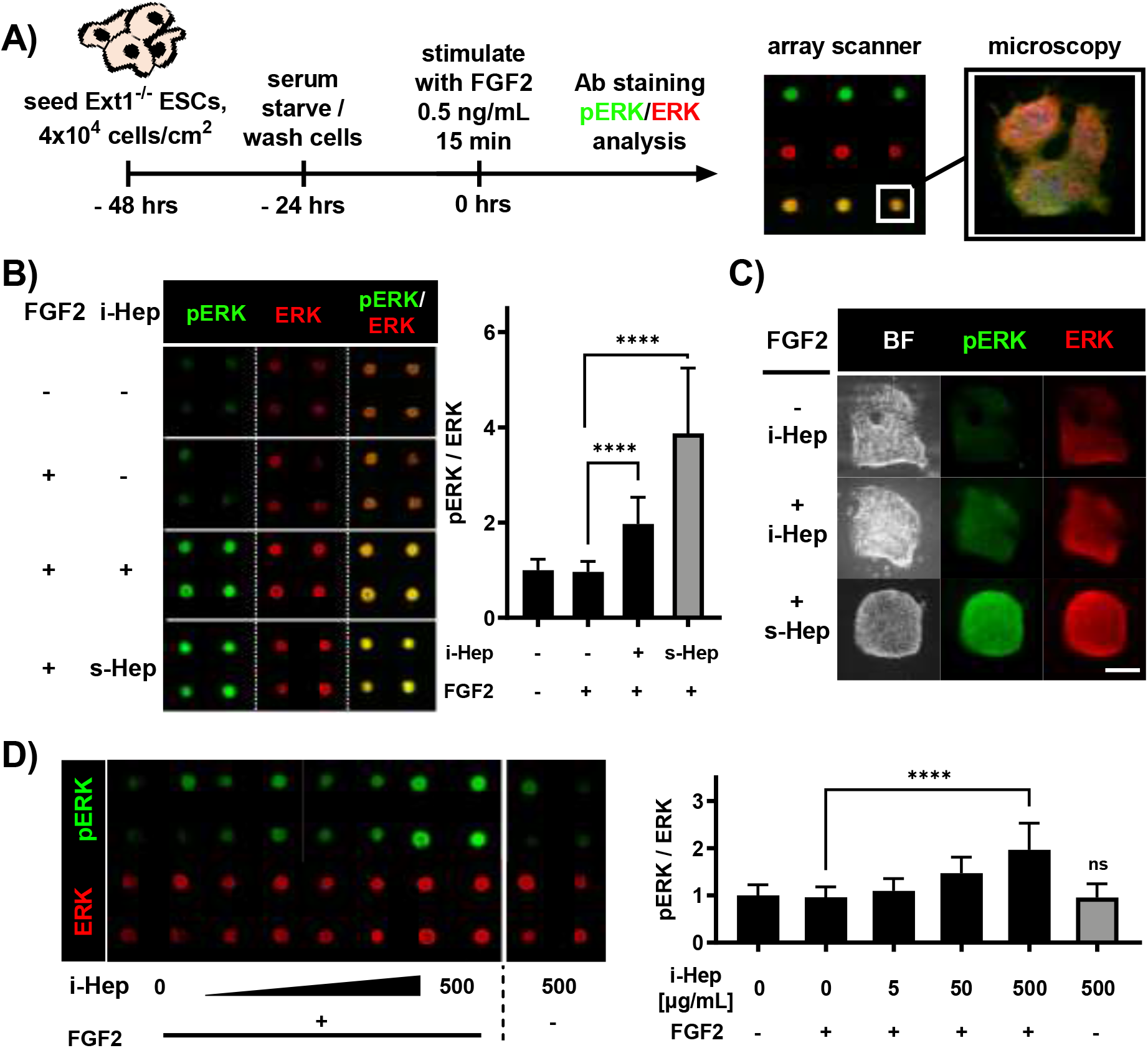
Analysis of FGF2 mediated MAPK pathway activation on a cellular heparin array. A) Timeline for mESC stimulation experiments on microarray. Cells were seeded and proliferated for 24 hours, washed and serum starvation for 24 hours, then stimulated with 0.5 ng/mL FGF2 for 15 mins, followed by fixation and immunodetection of pERK and ERK. B) Rapid assessment of cell signaling in the presence of immobilized heparin, using soluble heparin as a benchmark for MAPK activation. C) Microscopy of immunostained colonies showing pERK induction in i-Hep and robust response with s-Hep addition. Scale bar = 250μm. D) Stimulation with FGF2 shows on immobilized heparin elicits dose-dependent MAPK response with the addition of FGF2, showing tunability of cellular response in arrayed ECM microenvironments. (Bars represent mean values and standard deviation for at least 9 replicate colonies per condition)

## Conclusions

We have developed an array platform for direct, rapid and multiplexed profiling of extracellular glycosaminoglycan activity on growth factor signaling in live embryonic stem cells. The arrays were generated by printing and physisorption of GAG polysaccharides chemically crosslinked with extracellular matrix proteins onto polyacrylamide hydrogel substrates. The crosslinking facilitated the retention of the polysaccharides on the array during cell culture without altering the protein-binding specificity of the polysaccharides. The immobilized GAG structures were able to facilitate GF-mediated activation of signaling events in live embryonic stem cells which could be detected and quantified using immunofluorescence. This array offers a convenient platform to systematically analyze biological activities of extracellular GAGs in stem cell signaling and to accelerate the development of new bioactive glycomaterials for stem cell-based therapeutic applications by capitalizing on the rapidly expanding repertoire of available synthetic,^[16]^ chemoenzymatic,^[17,18]^ and recombinant glycosaminoglycan structures.^[19]^

## Acknowledgements

This work was supported in part by the NIH Director’s New Innovator Award (NICHD: 1DP2HD087954-01). K. G. is supported by the Alfred P. Sloan Foundation (FG-2017-9094) and the Research Corporation for Science Advancement via the Cottrell Scholar Award (grant # 24119)

