## supplemental information for "Stem Cell Microarrays for Assessing Growth Factor Signaling in Engineered Glycan Microenvironments"

### Supporting Information

**List of abbreviations**

APS- ammonium persulfate

DMEM- Dulbecco’s modified eagle medium

DMSO - dimethylsulfoxide

EDC- 1-Ethyl-3-(3-dimethylaminopropyl)carbodiimide

EtOH- ethanol

ERK- extracellular regulated kinase

EXT1- exostosin 1

FBS- fetal bovine serum

FGF2 – fibroblast growth factor 2

GF- growth factor

HABP – hyaluronic acid binding protein

HEPES- (4-(2-hydroxyethyl)-1-piperazineethanesulfonic acid)

KO-DMEM- knock Out Dulbecco’s modified eagle medium

LIF- leukemia inhibitory factor

MAPK- Mitogen activated protein kinase

MQ- Millipore milliQ purified water

NEAA- non-essential amino acids

NHS- N-hydroxysuccinimide

NMR- nuclear magnetic resonance

PBS- phosphate buffered saline

pERK- phospho-extracellular regulated kinase

TEMED- tetramethlyethlyenediamine

v/v – volume/volume

w/v – weight/volume

Table 1. Biological reagents and consumables

| Reagent | Supplier | Catalog # |
|---|---|---|
| 2-mercaptoethanol | Gibco | 21985-023 |
| DPBS | Corning | 21-031 |
| FBS | Gibco | A4766801 |
| FGF2 | Gibco | PHG0264 |
| Heparin | Acros organics | 411212500 |
| KO-DMEM | Gibco | 10829018 |
| L-Glutamine | Gibco | 25030-081 |
| LIF | Milipore ESGRO | ESG1107 |
| NEAA | Gibco | 11140-050 |

Table 2. Staining reagents and antibodies used for visualization

| Antibody/protein | Concentration | Catalog # |
|---|---|---|
| Rabbit Phospho-ERK1/2 | 1:200-1:300 | 4370 |
| Mouse Total-ERK1/2 | 1:200-1:500 | 4696 |
| Oct4 | 1:100 | Sc-25401 |
| Nestin | 1:250 | MAB353 |
| Anti-mouse-AF555 | 1:1000 | 13 |
| Anti-Rabbit AF647 | 1:1000 | 8 |
| CS56 | 1:100 | 11570 |

#### General methods, materials, and instrumentation

##### General chemistry

All chemicals, unless stated otherwise, were purchased from Sigma Aldrich. Purchased starting materials were used as received unless otherwise noted. Heparin and desulfated heparinoids were purchased from Iduron (Manchester, UK). The selectively desulfated heparinoids originated from the unmodified heparin used in this study. Iduron reported the average molecular weight of the parent heparin approximately 12,000 g/mol. Solvent compositions are reported on a volume/volume (v/v) basis unless otherwise noted.

##### General instrumentation

Nuclear magnetic resonance (NMR) spectra were collected on a Bruker 300MHz NMR spectrometer. Spectra are reported in parts per million (ppm) on the δ scale relative to the residual solvent as an internal standard. Brightfield images of live cells were taken using ZEISS Axio Observer microscope. Fixed cells were fluorescently imaged using a Keyence BZX-700 fluorescent microscope. Microarray slides used for protein binding assays were assessed using an Axon GenePix 4000B microarray scanner (Molecular Devices). All microarray experiments, cellular and protein binding, we performed using steel spring ProPlate gaskets (Gracebiolabs, cat. No. 248864) which were attached to the array slide.

**Preparation of NHS-activated glycosaminoglycans**

A HEPES buffer (100 mM, 15.0 mL, pH 7.4) was first prepared. Then, GAG (0.17 μmol) was dissolved in 200.0 μL of the HEPES buffer, 4 equiv. per GAG chain (0.68 μmol) of NHS was added via an aliquot from a stock NHS solution in HEPES and stirred overnight at 4°C to afford heparin NHS-ester. The activated NHS ester solution was diluted to a total volume of 2.5 mL using in MQ water, loaded onto a PD-10 column, and eluted with 2.5 mL MQ water. The solution was lyophilized to afford intermediate heparin NHS-ester (2.0 mg, 100% mass recovery). The resulting 2.0 mg of heparin-NHS product was dissolved in a PBS solution containing gelatin (10.0 mg/mL, 200 μL). The reaction was allowed to proceed overnight. The crosslinked glycosaminoglycan product was purified through a PD-10 column (2.5 mL loading volume, 2.5 mL elution volume). The resulting solution was lyophilized to afford purified gelatin containing crosslinked glycosaminoglycans as a white spongy solid (4.1 mg, 100% mass recovery based on BCA assay). The same stoichiometry was used for all glycosaminoglycan conjugates.

#### GAG array construction and validation

##### Glass slide cleaning

Untreated 25x75 mm glass microscope slides were loaded into a steel slide rack and submerged in a crystallization dish filled with MQ $H_2O$. The slide rack was washed five times with water, allowing the slides to remain in the last water wash for 30 minutes on a rocker. After 30 minutes, the water was removed and replaced with acetone. This solution rocked for 30 minutes, covered. The acetone was then removed and replaced with MeOH, was once more rocked for 30 minutes, covered. The MeOH was removed and replaced with a solution of 0.05 M NaOH (1g NaOH, 500mL $H_2O$), and rocked for 2 hours. Slides were then rinsed three times in MQ $H_2O$ and subsequently spin dried (500 rpm, 5 min). The slides were then lightly blow-dried using 0.22 μm filtered air. Once dried, slides can then be placed into a vacuum oven to dry at 70 °C, 20 PSI, and safely stored for up to a month.

##### Glass slide silanization

Dry, NaOH etched slides (in a steel rack) were placed into a solution of 2% 3-(trimethoxysilyl) propyl methacrylate 98% toluene (v/v), and rocked for 1 hour. The solution was then removed and the slides were washed three times in fresh toluene to remove residual 3-(trimethoxysilyl) propyl methacrylate. The slides were spin dried (500 rpm, 5 min), blow-dried with 0.22 μm filtered air, and placed in a desiccator overnight. The slides also can be dried in a vacuum oven to dry at 70 °C, 20 PSI for one hour, and can be stored for up to a month. The slides are then glutaraldehyde activated, by rocking the slides for 2 hours in a 0.05 % solution of glutaraldehyde in MQ $H_2O$. The slides were spin dried (500 rpm, 5 min), blow-dried with 0.22 µm filtered air, and placed in a desiccator overnight. The slides also can be dried in a vacuum oven to dry at 70 °C, 20 PSI for 15 minutes. Slides stored in this fashion can be used for up to one month.

**Deposition of acrylamide hydrogel on glass slides**

An aqueous 30% acrylamide solution was prepared by addition of 2.85 g acrylamide, 0.150 g (19:1 acrylamide/bisacrylamide) to 10.0 mL $H_2O$. A separate solution of 10% (w/v) ammonium persulfate (APS) solution was prepared in MQ $H_2O$. The polymerization solution was then prepared by combining 985 µL MQ $H_2O$, 500 µL 30% 19:1 Acrylamide solution, 15 µL of 10% APS, and 0.6 µL TEMED, in that order. Immediately following TEMED addition, 110 µL aliquots of the polymerization solution was placed in the center of glutaraldehyde activated methacrylate slide. Before polymerization occurs, a cover slip was quickly placed on each slide with a polymerization aliquot in such a way that no air bubbles form. To ensure the absence of air bubbles, the coverslip was angled at ~ 45° over the glass slide so that the liquid drop makes contact with both the slide and the coverslip. Then, the coverslip was lifted and widen the angle relative to the slide to pull the droplet towards the edge of the slide (where the vertex of the created angle is). When the droplet reaches the back edge of slide it will begin to widen and distribute along the length of the slide slightly, allowing the coverslip to be gently placed onto the slide, reducing the space and angle between the slide and coverslip until the coverslip can no longer be held without disturbing the slide, at which point it can be dropped. After the

TEMED addition, polymerization typically takes about 3-5 minutes. The polymerization was then allowed to proceed for 2 hours, and the slides were loaded into a steel slide rack with the coverslips still on, and allowed to sit in $H_2O$ for 15 min, causing the coverslip to loosen on the slide as the hydrogel expands. Slides were removed from water, and using a razorblade, the coverslips were gently removed. The hydrogel exposed slides were then carefully reloaded into the steel slide racks and submerged in a crystallization dish filled with $H_2O$. The $H_2O$ was replaced every 24 hours for a total of 48 hours of washing. After the 48-hour wash, the slides were spin dried and placed hydrogel-side-up onto a slide warmer heated to 50 °C for 10 minutes, or until slides were partially dehydrated for storage.

**Printing of GAG arrays**

Microarrays were printed using an SpotBot Extreme microarrayer (ArrayIt). Arrays were printed in 65% humidity using 500 µm spot pins (SMP15). While the number of spots varies from array to array, spacing between spots was consistently 1400 µm in arrays used for cellular culture. For protein binding assays, spots were spaced 750 µm apart, as the extra space was not required for to accommodate cellular growth. When designing the spot layout, the print parameter option MAUI4 was selected, and the lateral and vertical offset were 1 and 3 mm respectively. Arrays were always printed with 0.5 mg/mL porcine gelatin bloom 180 in PBS supplemented with 10% glycerol and 0.03% triton X-100. When concentration gradients we printed, the lowest concentration was always printed last. The gelatin or heparin crosslinked gelatin was printed at 500 µg/mL.

After printing, slides are placed into a slide holder and allowed to dry overnight at 4 °C. Prior to use, slides were washed in MQ $H_2O$ for 2 minutes by loading slides into a steel rack and rapidly and repeatedly dipping slides into a crystallization dish full of MQ $H_2O$. After this, slides were washed three times in PBS, 15 minutes, and then spin dried by centrifuging slides at 500 rpm for 5 minutes. Following this, the slides are snap dried by placing the cells array-side-up onto a slide warmer heated to 50 °C for 10 minutes, or until slides are clearly dried. At this point, slides could be stored or used for up to a month.

**Growth factor binding on arrays**

Microarrays used for protein binding assays were equipped with a 4-well gasket chamber and then blocked for 45 minutes with 250 μL of filtered PBS solution containing 1% BSA and 0.1% tween-20. After blocking, protein binding incubations were performed at 4 °C for 90 minutes in blocking solution with 10nM AF647-FGF2. Between protein incubations, wells were washed four times with blocking solution. After all incubations, a final series of three PBS washes for 15 minutes each were performed, the slide was spin dried, and scanned using a microarray scanner.

**Preparation of AF647-FGF2**

100 μg Human FGF2 was dissolved in 200 μL of HEPES buffer (200 mM, pH 8.4). Then, 20 μL of a 20 mg/mL heparin solution in MQ $H_2O$ was added to the FGF2 solution and 10 minutes were allowed to pass, at which point 2 μL of a 10 mg/mL NHS-AF647 solution in DMF was added to the solution. The reaction was gently rocked for 3 hours, and was quenched by the addition of 80 μL of a 20 mg/mL glycine solution in MQ $H_2O$. To purify

the reaction, a heparin sepharose column (1 mL) was prepared and used. To an empty column, 500 µL of heparin sepharose was added and centrifuged at 1000 rpm for 15 seconds to remove the liquid from the heparin sepharose. Next, 500 µL of elution buffer (3 M NaCl, 2% BSA, 20 mM HEPES, pH 7.4) was run through the column and removed by spinning 1000 rpm for 15 min. This is repeated once, and the column was rinsed twice more with wash buffer (0.5 M NaCl, 0.2% BSA, 20 mM HEPES, pH 7.4). The quenched reaction was loaded onto the column. Two separate columns were used to avoid overloading. Half of the reaction mix was loaded onto a 1 mL column and the column was spun at 1000 rpm for 15 seconds. Then the eluted solution was reloaded onto the column and spun again at the same speed and duration. Then, 500 µL of the wash buffer was loaded and spun through at 1000 rpm for 15 seconds. This was repeated five times, and then purified AF647-FGF2 was eluted with 500 µL of elution buffer. The elution was repeated with fresh buffer and the two fractions were combined with each other, as well as the elution from the column done in parallel, to bring the final volume to 2 mL of FGF2 protein. The resulting solution was 2.9 µM of FGF2.

#### Cell culture and biological assays

##### ESC culture

All mouse embryonic cell lines were cultured feeder free in treated plastic well plates at 5% $CO_2$ and 37 °C. Cells were cultured in ESC maintenance media consisting of KO-DMEM media supplemented with 10% fetal bovine serum, non-essential amino acids, L-glutamine, 2-mercaptoethanol and LIF. Serum free media is of identical composition to ESC maintenance media except for the exclusion of FBS. Cells were passaged every other day and spilt at a ratio of 1:10 ($10^5$ cells).

##### Sterilization of arrays

Arrays were sterilized for cell culture in a laminar flow tissue culture hood by placing arrays and autoclaved gaskets in 100% EtOH for 5 minutes, followed by a sterile PBS wash. The gasket was then assembled onto the slide, and the slide is washed with PBS and left under UV light for at least 15 minutes this is repeated twice, each time with fresh PBS.

##### Seeding arrays

At least 20 minutes before seeding, 500 μL of LIF containing ESC maintenance media was added to each well of a 4 well gasket attached to an acrylamide slide with an array printed upon it. Cells were then seeded onto the array in a volume of 1 mL, bringing the final volume to 1.5 mL 24 hours after seeding, the outlines of gelatin spots became noticeable due to cells growing upon the spots, and the slide was washed once with DMEM and the media appropriate for the desired experiment is added onto the plate in a 1 mL volume.

#### Growth factor stimulation

Cells were seeded onto microarray wells at a density of $4x10^4$ cells/$cm^2$ and allowed to adhere for 24 hours in ESC maintenance media. After this time, cells were washed with PBS once to removed unbound cells, and media was switched to serum free ESC media for the next day. Following serum starvation, cells were washed with DPBS and treated with serum free media containing various amounts of FGF2 with or without 5 µg/mL heparin. Immediately following starvation, cells were returned to the incubator for 15 mins. After this incubation period, cells were placed directly onto ice for immunocytochemistry.

#### Immunocytochemistry

After stimulation cells were immediately washed with cold DPBS and fixed for 10 minutes at room temperature in 4% paraformaldehyde. Then cells were washed 3x with cold PBS and cellular membranes were permeabilized using cold methanol for 20 minutes. Cells were then washed 3x with PBS and blocked for 1 hour at room temperature with immunocytochemistry blocking buffer (3% (w/v) BSA, 2% (v/v) goat serum). The appropriate primary antibody was applied overnight in ICC blocking buffer at 4 °C. Cells were washed 3x with PBS and corresponding secondary antibodies were applied for 1 hour at room temperature. Cells were washed 3x with PBS and nuclei were stained with hoescht for 15 minutes at room temperature. Cells were then washed 3x with PBS and mounted overnight at room temperature using ProLong Gold antifade (cell signaling, Cat. #9071). The next day, cells were subjected to fluorescent microscopy imaging or scanner analysis using an Axon GenePix 4000B microarray scanner (molecular devices), equipped with a Cy3 and Cy5 filter.

**Statistical Analysis**

All mathematical analyses were performed using GraphPad Prism 9.0. The statistical significance of a single comparison was performed using the built-in analysis (Student's *t* test). In general, each condition was conducted in duplicate in each experiment, and at least two independent biological replicates were used to derive conclusions. Bar graph values represent mean ± SD. Thresholds for significance for all tests is set as *, $p < .05$; **, $p < .01$; ***, $p < .001$; ****, $p < .0001$.

Figure S1. Optimization of passivation method for stem cell culture on arrays.

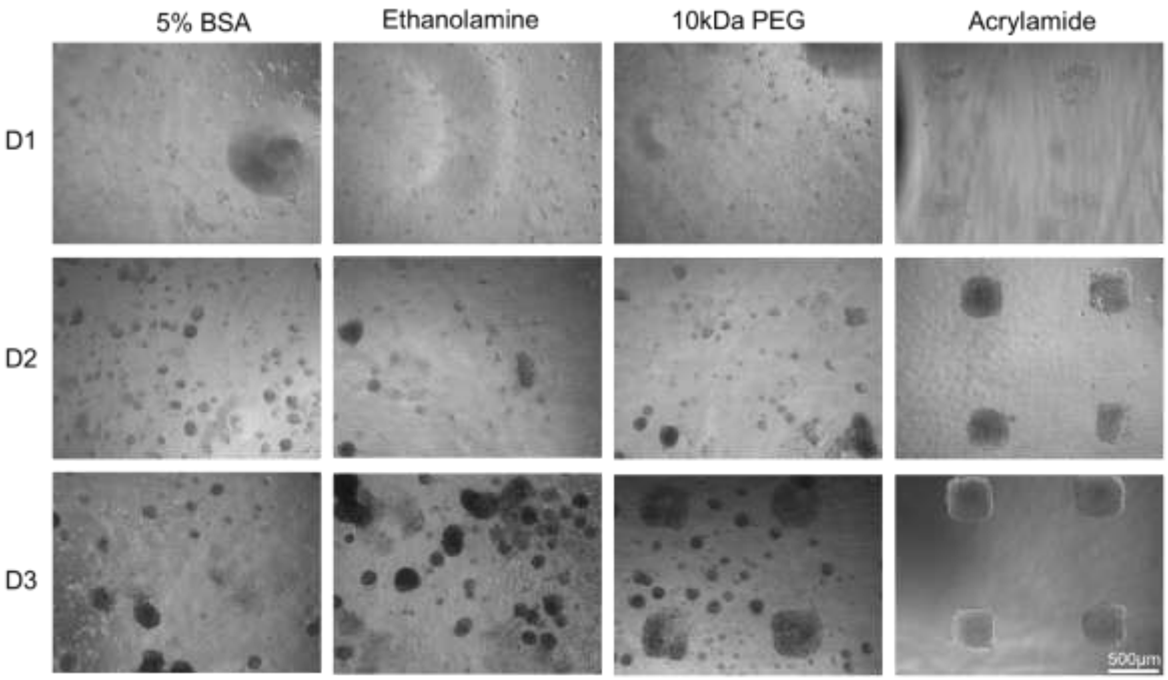

To generate a spatially segregated cellular microarray capable of multiplexed assays, several passivation strategies were tested. Epoxide functionalized slides (Thermofisher) were passivated using established passivation protocols[1,2], printed with 0.5 mg/mL gelatin in the absence of crosslinked GAGs, and tested by seeding 4x$10^4$ Ext1$^{-/-}$ ESCs and visually monitoring growth over three days using. For passivation, Slides were incubated at room temperature overnight with either 5 % Bovine Serum Albumin (BSA) solution, 100mM ethanolamine in pH 8.5 borate buffer. For 10kDa PEG, the slides were subjected to 10 mg/mL of 10 kDa amino-peg in PBS containing 62 mM $K_2SO_4$ overnight at 37 °C. Acrylamide slides were prepared as described, using established procedures. Acrylamide surface passivation afforded the most complete passivation of the surface with little unrestricted cellular growth. (D1 = Day 1, scale bar = 500 µm)

Figure S2. Quantification of Erk phosphorylation activity for *Ext1*$^{-/-}$ ESCs stimulated with FGF2 on arrays by fluorescence microscopy (extended data for Fig 4C).

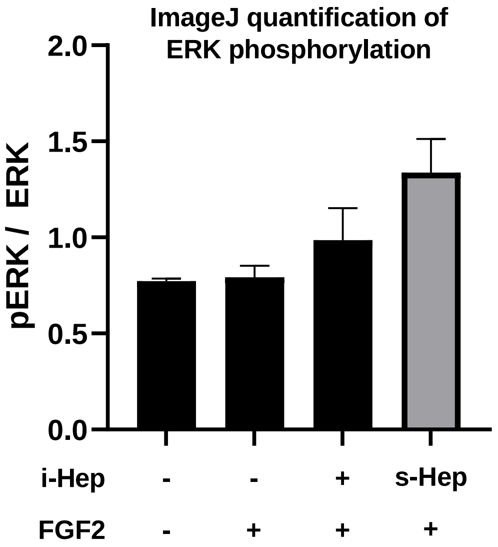

*Ext1*$^{-/-}$ ESCs were cultured on arrays with or without crosslinked heparin (500 µg/mL) were stimulated with FGF2 (0.5 ng/mL). Condition supplemented with soluble heparin (5 µg/mL) was used as a positive control. Immunoassayed colonies on the array were imaged and levels of p-ERK and ERK were quantitated using ImageJ software. Bars represents the means and standard deviations of at least two colonies for each condition.

Figure S3. Microarray images of arrayed *Ext1*$^{-/-}$ ESC colonies stimulated with FGF2 and immunoassayed for p-ERK and ERK (Extended data for Figure 4D).

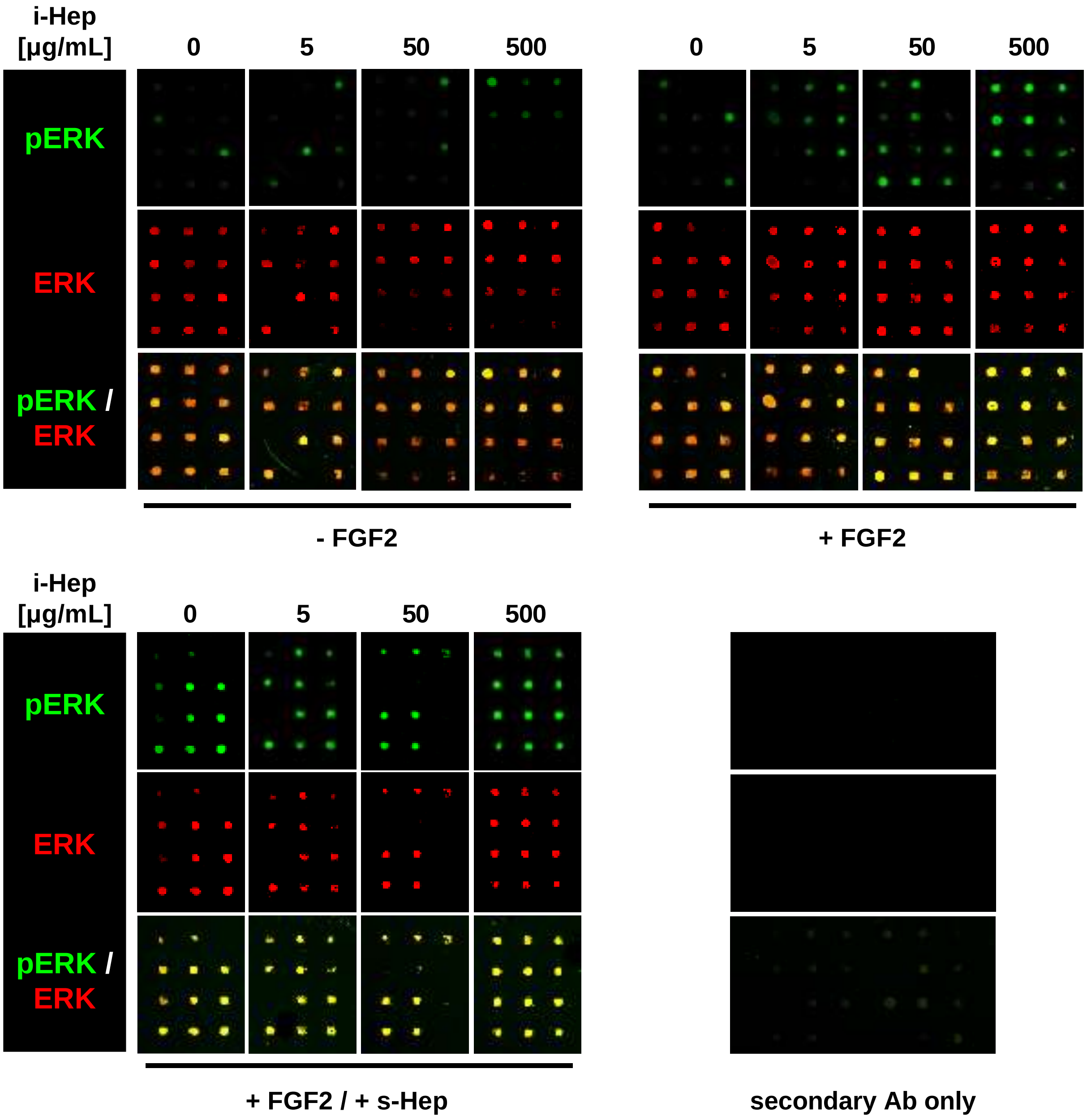
